# Revised Mechanism of Hydroxyurea Induced Cell Cycle Arrest and an Improved Alternative

**DOI:** 10.1101/2024.03.02.583010

**Authors:** Alisa E. Shaw, Jackson E. Whitted, Mattias N. Mihelich, Hannah J. Reitman, Adam J. Timmerman, Grant D. Schauer

## Abstract

Replication stress describes various types of endogenous and exogenous challenges to DNA replication in S-phase. Stress during this critical process results in helicase-polymerase decoupling at replication forks, triggering the S-phase checkpoint, which orchestrates global replication fork stalling and delayed entry into G2. The replication stressor most often used to induce the checkpoint response is hydroxyurea (HU), a chemotherapeutic agent. The primary mechanism of S-phase checkpoint activation by HU has thus far been considered to be a reduction of dNTP synthesis by inhibition of ribonucleotide reductase (RNR), leading to helicase-polymerase decoupling and subsequent activation of the checkpoint, mediated by the replisome associated effector kinase Mrc1. In contrast, we observe that HU causes cell cycle arrest in budding yeast independent of both the Mrc1-mediated replication checkpoint response and the Psk1-Mrc1 oxidative signaling pathway. We demonstrate a direct relationship between HU incubation and reactive oxygen species (ROS) production in yeast nuclei. We further observe that ROS strongly inhibits the *in vitro* polymerase activity of replicative polymerases (Pols), Pol α, Pol δ, and Pol ε, causing polymerase complex dissociation and subsequent loss of DNA substrate binding, likely through oxidation of their integral iron sulfur Fe-S clusters. Finally, we present “RNR-deg,” a genetically engineered alternative to HU in yeast with greatly increased specificity of RNR inhibition, allowing researchers to achieve fast, nontoxic, and more readily reversible checkpoint activation compared to HU, avoiding harmful ROS generation and associated downstream cellular effects that may confound interpretation of results.

## INTRODUCTION

DNA replication is plagued by a multitude of challenges, collectively known as replication stress, which threaten genomic stability and place cells at risk of mutagenesis and/or death. Physical barriers to normal replication in S-phase are numerous and include endogenous obstacles like DNA secondary structural elements and DNA-bound proteins, as well as DNA lesions directly caused by genotoxins including metabolic intermediates, exogenous chemicals and UV light, to name a few (1, 2). Another important source of replication stress includes the depletion of deoxynucleotide triphosphate (dNTP) pools, causing polymerase stalling during elongation— accordingly, dNTP levels are strictly regulated during S-phase (1, 3–6). One common thread in replication stress is the occurrence of helicase-polymerase uncoupling: the eukaryotic helicase CMG (Cdc45, Mcm2-7, GINS) encircles the leading strand and normally forms a stable complex with the leading strand Polymerase (Pol) ε (7, 8), but this complex rapidly separates upon Pol ε stalling (9, 10). This functional uncoupling of helicase and polymerase activity has been shown to be the primary activator of the S-phase replication checkpoint, a pathway that has evolved to stabilize replication forks in the presence of replication stress.

In the S-phase checkpoint response, Mec1, the primary replication checkpoint kinase in budding yeast (homologous to ATR in humans), is activated by stretches of RPA-coated single-stranded DNA resulting from runaway CMG helicase (1, 11–14). Activated Mec1 is recruited to replication forks to phosphorylate Mrc1, the Mediator of the Replication Checkpoint (Claspin in humans), a mediator kinase which travels with the replisome during elongation and which is required for both normal replication speed and checkpoint activation (15–22). When activated by the checkpoint, Mrc1 subsequently phosphorylates Rad53, which slows and stabilizes replication forks with the help of the Tof1/Csm3 pausing complex (Tipin/Timeless in humans) by phosphorylating CMG helicase and Mrc1 itself (23–28) (**Fig. 1a**). Mrc1 null yeast strains are viable but progress through S-phase slowly and demonstrate increased sensitivity to chemically induced replication stress (23). Interestingly, the roles of Mrc1 in replication speed and checkpoint signal transduction appear to be separable: yeast expressing only Mrc1^AQ^, a version of Mrc1 containing Ala mutations of all 17 SQ/TQ residues (a motif demonstrated to be the primary target of the Mec1 family (29)), were shown to be completely nonfunctional in S-phase checkpoint activation while maintaining a normal S-phase duration (15). Thus, yeast strains bearing *mrc1*^*AQ*^ can be used to study the effects of replication stress under surveillance of a disabled replication checkpoint apparatus with otherwise fully functional replication and cell cycle progression.

**Figure 1.**
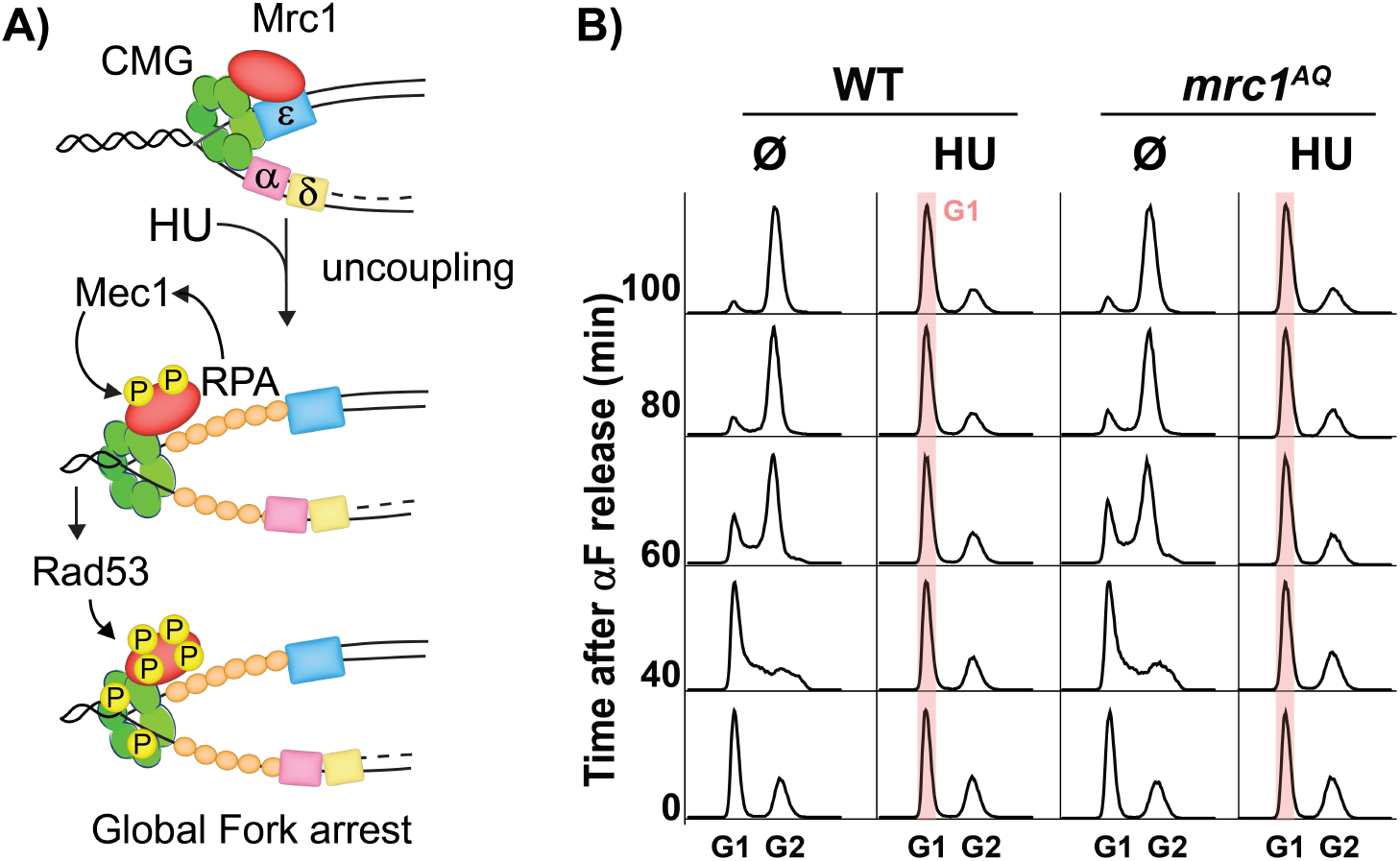
Hydroxyurea-induced cell cycle arrest without the S-phase checkpoint response. **(A)** Schematic of replication checkpoint. Uncoupling activates Mec1, which phosphorylates Mrc1, activating Rad53, which phosphorylates Mrc1 and CMG, stabilizing and arresting forks. **(B)** G1/S arrest independent of the Mec1/Mrc1 pathway. WT or *mrc1*^*AQ*^ cells synchronized in G1 by alpha factor (αF) were released into media containing either 0 or 0.2 M HU, and flow cytometry profiles were acquired at the indicated times. Red bar indicates arrest at the G1/S transition.

Hydroxyurea (HU) is an FDA-approved chemotherapeutic and is to date the most widely utilized drug to induce cell cycle arrest in the literature. HU usage is particularly prevalent in budding yeast, where concentrations up to 0.2 M are routinely used to achieve full cell cycle arrest. The primary mechanism of HU has so far been considered to be an inhibition of ribonucleotide reductase (RNR), the enzyme complex responsible for converting NDPs to dNDPs, which are required to produce dNTPs (30, 31). RNR is a heterotetramer comprised of a dimeric large catalytic subunit and a dimeric small regulatory subunit: Rnr1 is the main catalytic subunit, while Rnr3 is a nonessential paralog catalytic subunit; both these large subunits require the small subunits, Rnr2 and/or Rnr4 for activity (32). Inhibition of RNR by HU has been demonstrated to occur by electron transfer from HU to a catalytically requisite diferric tyrosyl radical center, thus limiting dNTP pools and stalling polymerases (31–35). However, there remain unexplained phenomena surrounding this proposed mechanism. For instance, it was observed that HU can still arrest replication forks in archaea which use a rare and separate RNR class that does not depend on this tyrosyl radical, without affecting dNTP pools (36). Further, it was shown in budding yeast, which cannot use nucleotide salvage pathways and thus strictly require RNR for dNTP supply, that HU only causes a moderate reduction of nuclear dNTP levels to concentrations that are readily utilized by DNA polymerases *in vitro* without reduction in polymerase activity, such that yeast would have to sense and respond to minor changes in dNTP levels through an as yet undescribed mechanism (37). A further complication to the usage of HU to induce replication arrest has been increasingly described in the literature, whereby HU results in the accumulation of reactive oxygen species (ROS) such as H_2_O_2_, possibly caused by radical chain reactions initiated by its hydroxylamine group (33, 38, 39). ROS can be teratogenic by directly damaging DNA, and DNA oxidation accumulates during HU administration, whereas these effects can be partially reversed by antioxidants (40–42). Besides being linked broadly to oxidative stress, many other potential targets of HU have been discovered, including catalase, carbonic anhydrase, and matrix metalloproteinases. It is indeed progressively more understood that ROS and its potential downstream targets is responsible for much if not most of the acute cytotoxicity of HU (33, 43, 44).

Increasingly, iron-sulfur (Fe-S) clusters have been discovered in various protein structures: to date over 200 Fe-S containing proteins have been described in humans, including many enzymes required for DNA metabolism. Fe-S clusters most frequently appear in the form of [4Fe-4S], a cubane-type arrangement of four Fe ions and four sulfide ions tightly coordinated by four negatively charged thiolate sulfurs of surrounding cysteine residues. Although the precise role(s) that these Fe-S clusters play are not fully understood for every protein, it is established that they are often enzymatic cofactors, can separately play roles in protein complex stability, and are broadly sensitive to the redox state of the cell (45–47). Intriguingly, it has been shown that primase regulatory proteins (Pri2 in yeast and p58 in human), part of the 4-subunit Pol α holoenzyme responsible for synthesizing RNA/DNA primers *de novo* during replication, possess a Fe-S cluster in their C-terminal domain (CTD) (48–53), and there is evidence that the Fe-S cluster in the human primase may act as a redox switch for DNA binding and primer handoff (54). It was demonstrated that all three eukaryotic replicative polymerases (Pols α, δ, and ε) contain an integral Fe-S cluster in their CTD, coordinated by a cysteine rich metal-binding motif called CysB (55) and an adjacent motif, called CysA, is thought to coordinate Zn^2+^ by default; although whether CysB in Pols α and Pol ε coordinate an Fe-S cluster or a Zn^2+^ metal remains unclear (46). An integral Fe-S cluster was recently observed in the N-terminal domain (NTD) of Pol2 (the polymerase subunit of Pol ε, the 4-subunit leading strand polymerase), coordinated by the CysX motif (56), and mutation of this domain was observed to abrogate polymerase and exonuclease activity by disruption of DNA binding (57). Further, disruption of the Fe-S coordination by CysB mutation in the CTD of Pol3 (the polymerase subunit of Pol δ, the 3-subunit lagging strand Pol) destabilized the complex and similarly compromised both its polymerase and exonuclease activities (55, 58). The effect on Pol δ was also observed *in vivo* by Vernis and colleagues, where mutation of its CTD CysB domain resulted in genome instability (59), corroborating earlier results from a genetic screen (60). This group has also demonstrated that HU activates the oxidative stress response pathway, which attenuates the deleterious oxidative effects of HU in part by upregulation of two critical genes involved in Fe-S biosynthesis, and additionally that H_2_O_2_ inhibits Leu1, an isopropylmalate isomerase containing a Fe-S cluster (61); these findings collectively led to speculation that various stress and/or anti-cancer drugs could be responsible for similar effects by oxidizing Fe-S clusters in cells (59, 61).

The present study was prompted by a surprising finding: HU-induced cell-cycle arrest in yeast independent of the replication checkpoint (see Results). In an effort to explain this confounding result, we wondered whether disruption of the various integral Fe-S clusters in eukaryotic Pol complexes could be responsible for the chemically induced replication arrest we observed in the absence of programmed stalling via Mec1/Mrc1 signaling. We therefore sought to quantify the extent to which HU induces ROS formation in yeast nuclei, testing the prediction that HU and/or H_2_O_2_ may partially inhibit the eukaryotic replicative Pols α, δ, and ε. During our attempt to understand the off-target effects of HU, we also developed a superior system in budding yeast called “RNR-deg” that directly targets the RNR apparatus with greatly improved specificity.

## RESULTS

### Cell cycle arrest by hydroxyurea independent of the S-phase checkpoint and Psk1 signaling

We initially constructed a *mrc1*^*AQ*^ strain, which is defective in checkpoint signaling and is less sensitive to HU compared to *mrc1◿*, but unlike *mrc1◿* still exhibits timely progression through S-phase (15). Since *mrc1*^*AQ*^ is unable to activate Rad53 and stabilize replication forks during HU-induced helicase-polymerase uncoupling (15), our working hypothesis was that *mrc1*^*AQ*^ would be unable to stall in the presence of HU as it lacks the ability to orchestrate programmed global fork arrest via the replication checkpoint (**Fig. 1a**). To test this, we synchronized WT or *mrc1*^*AQ*^ cells in G1 with alpha factor (αF) and released them into media containing 0 or 200 mM HU, assessing cell cycle state by flow cytometry. Surprisingly, we discovered that cells exposed to HU were arrested at the G1/S interface (**Fig. 1b**; scatter plots for this experiment and representative gating criteria used for all cytometry studies are shown in **Fig. S1**). Duch et al. recently reported that cells were permissive through S-phase when released from G1 into 50 mM HU (62), so we titrated HU into asynchronous WT or *mrc1*^*AQ*^ cells growing in YPD to see whether the stalling we observed was concentration dependent. We observed reasonable agreement with their data at 50 mM HU, the maximum used in their study: cells appeared to be at least partially permissive through S-phase. However, at higher HU concentrations, we observed a distinct accumulation of cells at the G1/S interface (**Fig. S2**), indicating an S-phase independent arrest caused by incubation with more than 50 mM HU. It was recently discovered that Psk1 kinase, having been identified in the oxidative stress response (63), is the primary kinase that targets Mrc1 in response to oxidative stress, at separate sites from those targeted by Mec1, resulting in delayed progression through S-phase under oxidative conditions (62). We therefore wondered whether Psk1 could be responsible for signaling to Mrc1 under oxidative stress and activating the checkpoint. We hypothesized that knocking out *psk1* might release the HU-dependent stalling we observed in the absence of Mec1/Mrc1 signaling. To the contrary, cells with a *psk1*◿ *mrc1*^*AQ*^ background released into 200 mM HU also stalled in G1/S in the presence of HU, indicating that the pausing we observed was entirely independent from Mec1/ATR signaling, but also from any oxidative stress response transduced via Mrc1 (**Fig. S3**). To further corroborate this, we compared these results to a *mrc1◿* strain. In agreement with previous reports (15, 23), we observed slow S-phase progression in *mrc1◿* cells compared to WT. Importantly, however, we also observed that G1-arrested *mrc1◿* cells released into media containing 200 mM HU displayed arrest at the G1/S interface, similar to the effect seen in the *mrc1*^*AQ*^ background, further supporting the finding that the S-phase checkpoint is not required for cell cycle arrest in the presence of high HU concentrations (**Fig. S4**). Taken together, these results indicate that HU can arrest cells in their cell cycle independent of the Mec1/Mrc1 and/or Psk1/Mrc1 pathways, and thus this drug likely acts outside of the S-phase checkpoint.

### A direct relationship between HU concentration and ROS generation in yeast cell nuclei

Since it has been shown that HU generates ROS in yeast, we wanted to query the extent to which ROS generation might be responsible for stalling cells. We detected nuclear ROS using dihydroethidium (DHE), an indicator that is converted to ethidium in the presence of ROS, consequently intercalating with DNA and exhibiting a strong red shift in fluorescence emission from 420 nm to 610 nm (see **Fig. 2a**). We grew yeast in log phase for 1 hour in the presence or absence of 200 mM HU, treated cultures with DHE for 1 hour, washed cells with PBS, and then examined fluorescence in cells by flow cytometry. In agreement with another study (61), we show a dramatically increased ROS signal from yeast cell nuclei in the presence of HU (**Fig. 2a**). To accurately assess the relationship between the amount of HU and ROS produced in yeast cells, we switched to a plate assay to quantify our results more robustly. As expected, we saw a hyperbolic increase in the fold change of red-shifted fluorescence in yeast nuclei with increasing amounts of H_2_O_2_ (the most common cellular ROS), likely plateauing due to a saturation of the conversion of DHE to ethidium; we also observed a strikingly similar trend when titrating HU into yeast (**Fig. 2b**). When directly correlating these values, we quantified a direct relationship between the fold change in fluorescence intensity caused by increasing [HU] versus that seen by increasing [H_2_O_2_] (*r* = 0.85); interpreting these curves within their quasi-linear ranges, it can be inferred that incubation with 0.2 M HU results ∼20 mM H_2_O_2_ at minimum in the nucleus, with an indeterminate upper bound due to signal saturation. Although DHE is an established reporter of oxidation by ROS like H_2_O_2_ and superoxide, it also reacts to nonspecific oxidation (64). We wanted to rule out the possibility that HU was causing red shifted DHE fluorescence directly, so we performed an *in vitro* assay containing DHE and plasmid DNA, in the presence of HU or H_2_O_2_. Critically, we observed that ROS activated red shifted fluorescence emission of the stain, while HU did not directly cause DHE to become fluorescent (**Fig. S5**). Considering this, and since HU was previously observed to inhibit Leu1 (an unrelated cytosolic enzyme containing a Fe-S cluster) *in vivo* but not *in vitro* (61), we conclude as others have that HU must be generating radical species through one or more cellular metabolic intermediates, possibly initiated by its hydroxylamine group (38, 61). Taken together, these data illustrate a direct relationship between the amount of HU cells are incubated with and the amount of ROS consequently generated in the nucleus.

**Figure 2.**
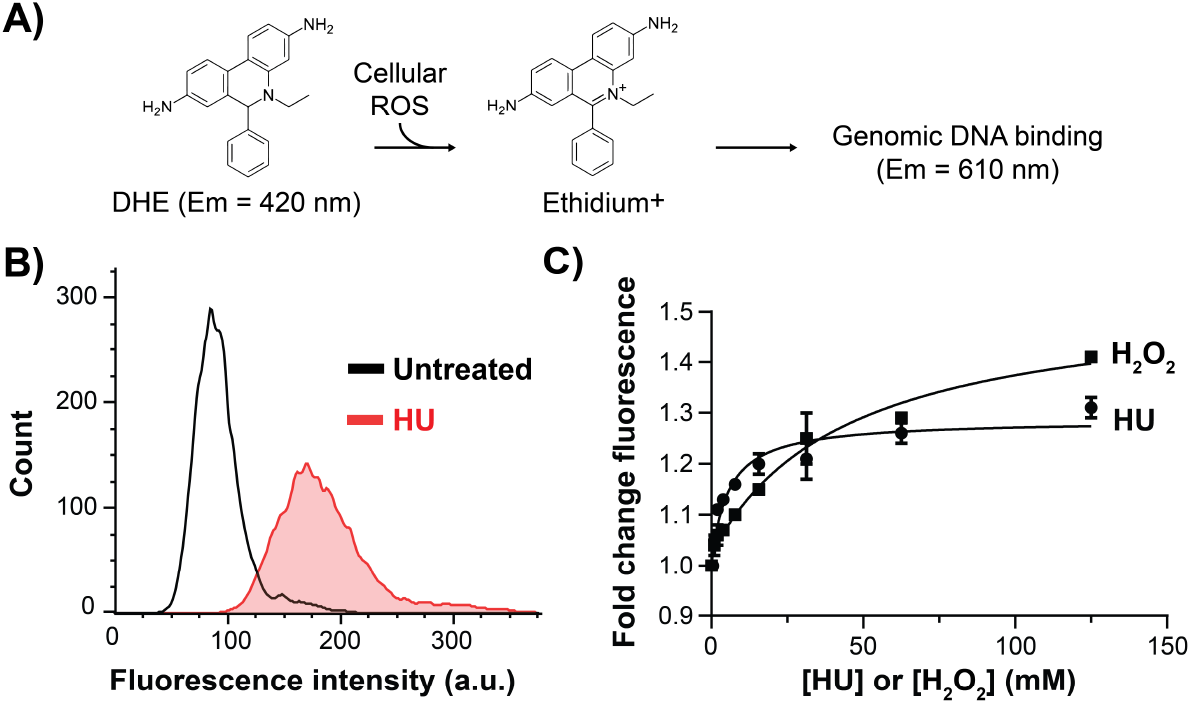
Direct relationship between HU incubation and ROS production in yeast cell nuclei. **(A)** Schematic of DHE assay. H_2_O_2_ and other cellular ROS oxidize DHE, converting it to ethidium, which intercalates nuclear DNA and subsequently exhibits red shifted fluorescence. **(B)** Cytometry profiles of red-shifted fluorescence intensity from DHE-stained cells in the absence or presence of 0.2 M HU. HU itself does not oxidize DHE (see Fig. S5). **(C)** Dose-response curves of yeast treated with increasing amounts of H_2_O_2_ (squares) or HU (circles); nonlinear curve fit R^2^ values were 0.94 and 0.97, respectively. Error bars represent STD from 3 replicates. Pearson’s *r* = 0.85 between both curves.

### Strong inhibition of eukaryotic replicative polymerases by ROS

Having observed cell cycle inhibition by HU irrespective of S-phase checkpoint activation, we wondered if either HU or the abundant ROS generated by HU might be responsible for inhibiting enzymes that were critical to cell cycle progression, like the three polymerases responsible for copying DNA: Pols α, δ, and ε. Performing *in vitro* strand-extension assays that utilize a radiolabeled primer annealed to an oligonucleotide template to monitor DNA replication, we first asked whether HU can directly inhibit DNA polymerases. In these assays, we loaded all substrates with PCNA sliding clamp using the RFC clamp loader and coated the template with RPA, an ssDNA binding protein. Pol δ requires PCNA clamp for activity, Pol ε can use clamp but does not require it, and Pol α does not work with or require clamp, however, we wanted to present all polymerases with identical reaction conditions for a more direct comparison. We first observed that DNA replication activity of both Pol δ and Pol ε was completely unaffected by the presence of up to 200 mM HU (**Fig. S6a**). Because we observed that HU generates ROS in yeast nuclei, we next asked whether H_2_O_2_ could inhibit DNA Pols α, δ, and ε. Indeed, we saw that all three polymerases were strongly inhibited by concentrations as low as 10 mM H_2_O_2_. We compared these results with Klenow Pol, a commercially available proteolytic product of *E. coli* DNA polymerase that possesses full polymerase and 3’-5’ exonuclease activity, and, like all prokaryotic polymerases, also lacks an integral Fe-S cluster. Importantly, we did not observe inhibition of Klenow at 50 mM H_2_O_2_, five times the amount that was inhibitory for the yeast polymerase complexes (**Fig. 3a**; example gel shown in **Fig. S6b**). Since our earlier results indicate that incubation with 200 mM HU is expected to yield roughly 20 mM H_2_O_2_ or more in cell nuclei, we conclude that replicative Pols are likely to be directly inhibited by the ROS generated by the high HU concentrations routinely used to arrest cells.

**Figure 3.**
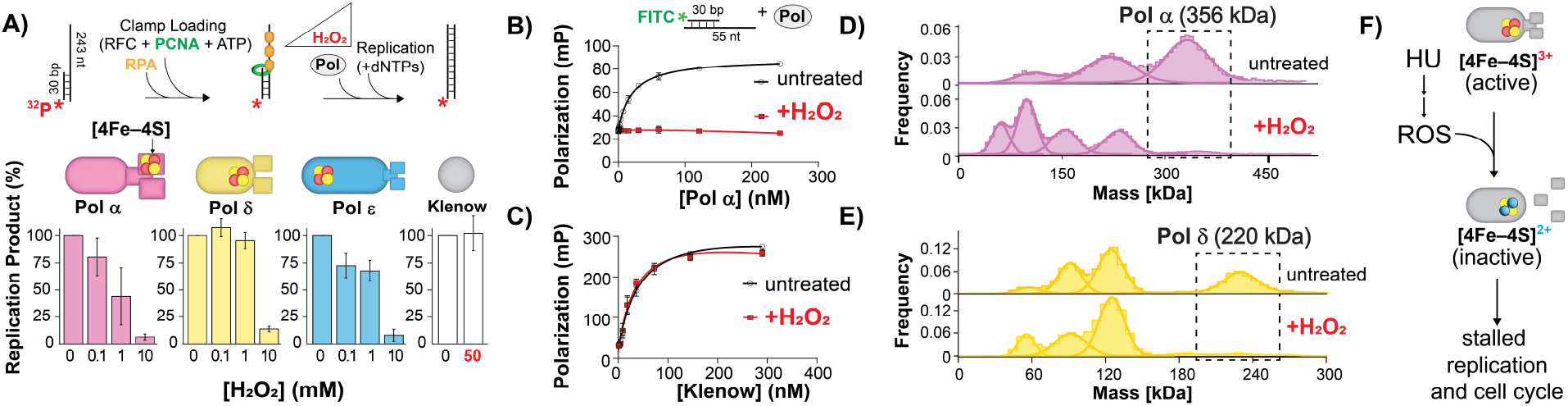
ROS inhibits the activity and DNA binding of eukaryotic polymerases by oxidizing their Fe-S clusters and destabilizing the complexes. **(A)** DNA strand extension activity of Pol α, Pol δ, or Pol ε (which contain Fe-S clusters) or Klenow Pol (which does not) was measured on clamp loaded, RPA coated template/primer substrates in the presence of the indicated H_2_O_2_ concentrations (top: assay schematic; Middle: Fe-S clusters are shown in Pri2 for Pol α, the CTD of Pol3 for Pol δ, and the NTD of Pol2 for Pol ε. Bar graphs are quantifications of three separate experiments; error bars represent STD (example gels in Fig S6). DNA template/primer binding activity of **(B)** Pol α or **(C)** Klenow Pol in the presence or absence of 50 mM H_2_O_2_ was measured by fluorescence polarization (top: assay schematic). Mass photometry histograms of **(D)** Pol α and **(E)** Pol δ are shown in the presence or absence of 9 mM H_2_O_2_. Dotted rectangle denotes the expected MW of fully assembled complex. **(F)** Model of cell cycle arrest by HU independent of RNR inhibition or checkpoint response. HU generates nuclear ROS, which oxidizes integral Fe-S clusters in replicative polymerases, causing polymerase complex dissociation and/or structural changes, halting global replication and stalling the cell cycle (see Discussion).

### DNA template/primer binding by Pol α is inhibited by ROS

We next asked whether the inhibition of polymerase activity via Fe-S cluster oxidation that we saw was localized to the polymerase active site or could be traced to other properties like DNA substrate binding, as was recently demonstrated with Pol ε (57). Pol α is responsible for synthesizing the hybrid RNA/DNA primers required for DNA replication and can also substitute for Pol ε and/or Pol δ (65), making it a good candidate as a central link to the cell cycle. We measured equilibrium binding of Pol α and Klenow Pol to DNA substrates by recording changes in fluorescence polarization of a fluorescein-labeled template/primer substrate and increasing amounts of Pol α. To force Pol α into the polymerase competent mode of DNA strand extension, the primer was dideoxy terminated and reactions included the next cognate dNTP (dTTP). Both Pol α and Klenow Pol bound this substrate with reasonably high affinity (*K*d=21.0 nM and 45.6 nM, respectively). Strikingly, the presence of 50 mM H_2_O_2_ resulted in complete abolishment of the DNA binding of Pol α (**Fig. 3b**) while this amount of H_2_O_2_ did not measurably alter substrate binding of Klenow Pol (**Fig. 3c**). Because the DNA binding clefts of Pol α and Klenow Pol are fundamentally similar, we attribute this difference to the existence of a labile, oxidation-sensitive Fe-S cluster in Pol α, but not in Klenow Pol. We next wanted to assess whether inhibition of Pol α DNA binding by H_2_O_2_ was due to a change in DNA binding affinity or by a precipitation or unfolding of the enzyme, so we measured the effect of increasing NaCl levels on the Pol α:DNA interaction in the presence or absence of H_2_O_2_. Without H_2_O_2,_ salt titration affected the result as expected by reducing both apparent *K*_d_ and by reducing the amount of apparent binding saturation. A similar effect could be seen in the presence of H_2_O_2,_ albeit at a reduced dynamic range, suggesting that Pol α remains soluble in the presence of H_2_O_2_ as evidenced by apparent but weak equilibrium binding that can be screened by salt (**Fig. S7**). At this point, however, we were unsure whether oxidation of the Fe-S cluster was responsible for disruption of local protein:DNA interactions or of the global structure of Pol α.

### ROS-induced dissociation of polymerase complexes with integral Fe-S clusters

Based on reports that Fe-S cluster oxidation may lead to polymerase and/or primase complex dissociation (55, 36, 58), as well as observations we have made during purification of Pols δ and ε in which oxidative conditions lead to complex dissociation, we hypothesized that disruption of the Fe-S cluster may lead to disassembly of polymerase holoenzymes. Using mass photometry to test this, we discovered that the Pol α and Pol δ complexes almost fully dissociated in the presence of >3 mM H_2_O_2_ (**Fig. 3d,e**; see **Fig. S8** for mass photometry histograms of Pol α and Pol δ acquired at further H_2_O_2_ concentrations and for a quantification of Pol α dissociation during H_2_O_2_ titration). The Pol ε complex was also observed to dissociate in the presence of H_2_O_2_ but was considerably less sensitive to H_2_O_2_ than Pols α or δ, remaining largely intact even at 45 mM H_2_O_2_ (**Fig. S8**). The MW of all Pol subunits are detailed in Fig S8. Collectively, our results indicate that ROS causes disassembly of polymerase complexes into their soluble subunits, inhibiting both DNA binding and polymerase activity (**Fig. 3f**).

### CysA and CysB of POL1 are dispensable for Pol α activity *in vivo*

Having observed dissociation of Pol α into its soluble subunits, we were interested to find that Pol α was so highly sensitive to H_2_O_2_ in terms of DNA binding and activity, particularly since our reaction scheme forced Pol α binding to DNA in a polymerase competent mode, i.e., not in its primase mode, implying either a direct modulation of the polymerase subunit (Pol1) and not the primase subunits (Pri1/Pri2) or a reliance on the primase domain for Pol1 binding to DNA. Pol α contains conserved CysA and CysB motifs in the CTD of its polymerase domain, but it remains unclear whether they coordinate Fe-S clusters (46, 55). This uncertainty is in large part due to the difficulty of purifying these complexes while retaining labile cofactors, however, genetics studies like those performed for Pol δ (59, 60) and Pol ε (56) can be informative regarding the importance of these metal binding sites. Pol α is essential, making it highly sensitive to changes in activity. To test whether the activity of Pol1 requires the coordination of Fe-S and/or Zn^2+^ by CysA and/or CysB, we first decided to degrade endogenous Pol1 using a C-terminal auxin-inducible degron (AID) tag (66) on POL1, which was lethal in the presence of the plant hormone 1-Naphthaleneacetic Acid (NAA) and the ubiquitin ligase OsTIR1 as determined by spot assays. We were then able to rescue this degradation of endogenous Pol1-AID with integrated an integrated copy of Pol1 under Gal control. Surprisingly, given the previously observed importance of CysB to Pol δ function, we were also able to rescue lethality by expressing Pol1 that had CysB and/or CysA metal binding motifs mutated by either Ser or Ala substitutions of the cysteines. At higher temperature (37°C), the Ala mutations of the CysB motif were unable to rescue degradation of endogenous POL1, which we consider likely to be a misfolding phenotype (**Fig. S9**). As such, these data indicate that Pol α does not require its CysA or CysB motifs for primase and/or polymerase activity *in vivo*. Based on this, we consider it likely that the Pri1/Pri2 domain, in which Pri2 contains an Fe-S cluster, is largely responsible for DNA template binding during DNA elongation (see Discussion).

### RNR-deg: a superior alternative to hydroxyurea-induced cell cycle arrest

These studies demonstrated that the oxidative toxicity of HU was high enough—and the specificity against RNR low enough—to warrant a new approach to cell cycle arrest, so we decided to directly target the RNR complex. Using an N-terminal AID degron tag on Rnr1 combined with an Rnr3 knockout, hereafter called “RNR-deg” (**Fig. 4a**), we observed lethality in the presence of both OsTIR1 and NAA by survival spot assays (**Fig. 4b**). Assessing by western blot, we observed that Rnr1 was rapidly and thoroughly degraded after 30 minutes (**Fig. S10**). We note that a C-terminal degron tag on Rnr1 was not viable even in the absence of NAA and OsTIR1 (not shown), likely because it prevented oligomerization, and that a strain bearing an N-terminal degron tag on Rnr2 was unaffected by the addition of OsTIR1 and NAA (see Fig. 4b). Importantly, when we synchronized cells in G1 and released them into media containing HU or NAA, we saw that RNR-deg arrests cells at G1/S in a manner than is indistinguishable from that of HU, using 3 orders of magnitude less NAA than HU (**Fig. 4c**). We observed a clear dose-response curve of the percentage of cells arrested in G1/S upon titration from 0 to 250 µM NAA **(Fig. S11a)**. Scatter plots from this experiment are shown in **Fig. S11b**, demonstrating that RNR-deg closely resembles HU treatment and does not significantly change cell shape. Treating with NAA at various timepoints after release from G1, we also demonstrate that RNR-deg can be used as a robust method of arresting cells in S-phase (**Fig. S12**). Critically, RNR-deg generates significantly less ROS in yeast nuclei than HU (**Fig. 4d**; compare RNR-deg signal to the HU signal in Fig. 2a). We note that a specific shutdown of dNTP production likely reduces the metabolic ROS load in cells, as we observed RNR-deg to have a decreased ROS signal compared to untreated cells (Fig. 4d). Further, we show that RNR-deg is more quickly reversible than HU; cells washed after a 4 hour treatment with RNR-deg reached G2 after 90 min, comparable to the speed of untreated cells, while cells washed after HU treatment were slower and had only partially reached G2 by 3 hours post wash (**Fig. S13**). This difference may reflect the relative inefficiency in removing three orders of magnitude more HU than NAA from cells (including the ROS that it generated) and could also be related to the time required to reassemble and/or reload polymerases after HU treatment. Taken together, we demonstrate that RNR-deg is a superior alternative to HU resulting in robust, rapidly reversible cell cycle arrest without the unnecessary production of harmful ROS.

**Figure 4.**
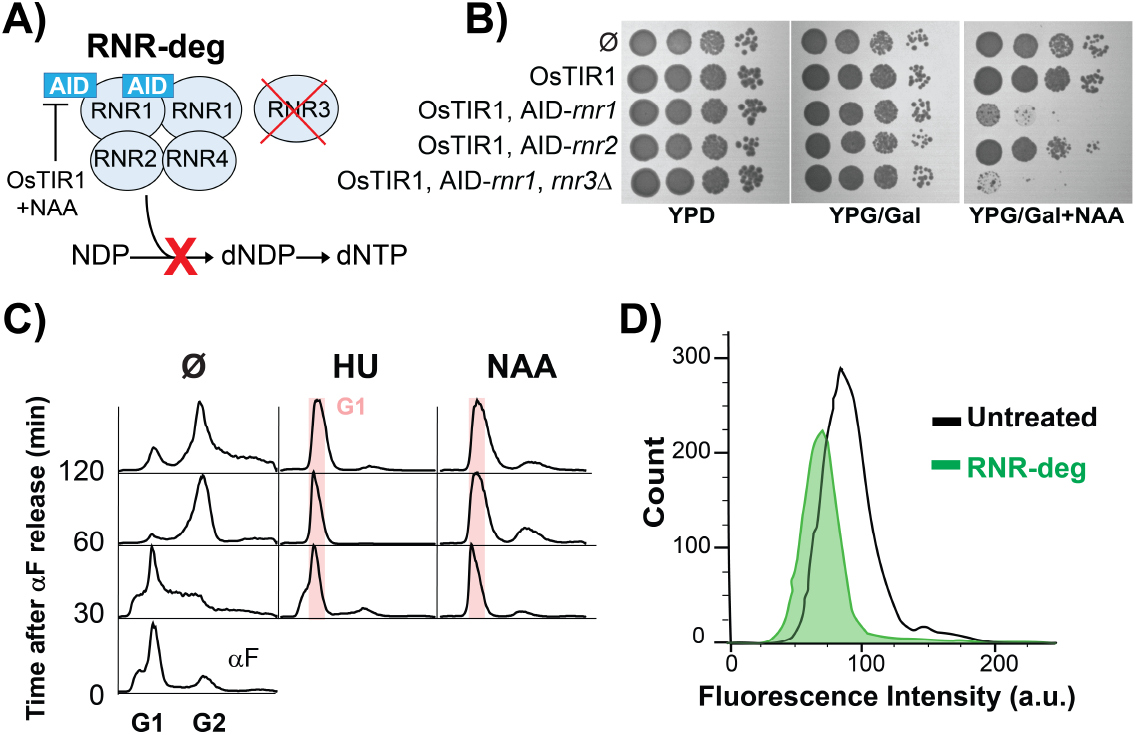
RNR-deg is a superior alternative to HU for inducing cell cycle arrest. **(A)** Schematic of RNR-deg. AID-Rnr1 is degraded by OsTIR1 in the presence of NAA, in an *rnr3◿* background, halting dNTP production required for DNA replication. **(B)** Survival spot assay of strains discussed in Results. RNR-deg is OsTIR1, AID-*rnr1, rnr3◿*; OsTIR1 expression requires Gal (bottom row). Spots contained 5×10^4^, 5×10^3^, 500, or 50 cells. **(C)** Flow cytometry profiles of RNR-deg cells that were synchronized in G1 by αF and released into YPG/Gal with the indicated treatment (200 mM HU or 250 µM NAA). **(D)** Cytometry profiles of red-shifted fluorescence intensity from DHE-stained RNR-deg cells in the absence or presence of 250 µM NAA; the untreated profile from Fig. 2b. is used for comparison.

## METHODS

### Strain construction

Our base strain, OY01, is derived from W303: *ade2-1 ura3-1 his3-11*,*15 trp1-1 leu2-3*,*112 can1-100 bar1Δ MATa pep4::KANMX6*) (67). The RNR-deg strain (AES38) was created by integrating Gal/OsTIR1 into the Ura3 gene into OY01. A miniAID1 tag (68) was added to the N-term of endogenous *RNR1* using CRISPR/Cas9. *RNR3* was replaced by a HygR cassette. The remaining strains used in this study (*mrc1*^*AQ*^, *mrc1*^*AQ*^ *psk1◿, mrc1◿*, deg-*POL1* and various *POL1* rescue constructs) were created using standard replacement of genes with various marker cassettes. Yeast cultures were grown in YPG/Gal media at 30°C unless otherwise stated. Further details for all strains can be found SI Materials and Methods.

### Flow cytometry

Cytometry experiments followed previously described methodology using Sytox Green to monitor DNA content of living cells (69) and were performed on a Cytek Aurora spectral cytometer. Reference samples of Sytox stained and unstained cells were performed for each experiment for live unmixing. Data was analyzed with FlowJo software using optimized gates that were static across each experiment, after first excluding small debris and large aggregates using FSC vs SSC plots and excluding non-DNA-containing debris with FSC vs. Sytox plots.

### ROS detection assay

All DHE experiments in yeast were performed with the RNR-deg strain, AES38. Cells were stained with 25 µg/mL DHE for 1 hour, washed, resuspended, and diluted 10-fold in PBS, and measured by spectral flow cytometry or in a 384-well plate reader (BMG Clariostar). Fluorescence intensity of red-shifted probe was measured as Ex: 488 nm and Em: 584 nm for cytometry experiments and Ex 360/40: Em: 600/40 in the plate assay. *In vitro* experiments in Fig S5 were 25 µL/well and were performed in the presence 1 ng/µL of supercoiled pAS1 plasmid DNA.

### Protein purification

The purification of Pol α, Pol δ, Pol ε, RFC, PCNA, and RPA has been previously described (65, 67, 70). The purity of protein complexes was assessed by coomassie stained SDS-PAGE, and all enzymes were assessed for activity *in vitro* before use in assays.

### Strand extension assays

Reaction volumes were 25 µL and contained 1.5 nM of the 85DS substrate, a 243 nt template primed with a 5’-^32^P-primer (construction described in Supplementary Methods). Reactions contained 25 mM Tris-OAc pH 7.5, 5% (vol/vol) glycerol, 50 μg/mL BSA, 2 mM TCEP, 3 mM DTT, 10 mM Mg-OAc, 50 mM K glutamate, 0.1 mM EDTA and were carried out at 30°C. DNA was first loaded with 20 nM RPA for 1 min, followed by 40 nM PCNA with 2 nM RFC and 1 mM ATP for 5 min. Pol α, δ, and ε were each added at 10 nM in the presence of the indicated HU and/or H_2_O_2_ concentrations. Reactions were initiated by the addition of 60 mM dNTPs, allowed to proceed for 5 minutes, and stopped with a STOP solution containing 1% SDS and 40 mM EDTA. Reactions were run by 7.5% denaturing PAGE. Gels were backed with Whatman 3MM filter paper, exposed to a storage phosphor screen for 6 hours, and visualized on a Typhoon FLA 9500 PhosphorImager (GE).

### DNA binding assay

Fluorescence polarization reactions were 20 μL and contained 25 mM Tris-OAc pH 7.5, 5% (vol/vol) glycerol, 50 μg/mL BSA, 2 mM TCEP, 3 mM DTT, 10 mM Mg-OAc, 125 mM NaCl (unless otherwise noted), 0.1% Tween-20, and 3 nM probe template/primer DNA, where the primer contains a 5’-fluorescein and is 3’ dideoxy terminated, in the absence or presence of 50 mM H_2_O_2_. Pol α titrations were scanned on a 384-well plate reader equipped with polarizing filters (BMG Clariostar; Ex: 482/16 nm, Em: 530/40 nm). Polarization is expressed in mP (P×10^-3^), where P = (I_VV_-I_VH_)/(Ivv+I_VH_); I_VV_ is the fluorescence intensity with vertical Ex/Em polarizers and I_VH_ is the fluorescence intensity with vertical and horizontal Ex/Em polarizers. Binding curves were fit to a single-site binding equation using GraphPad Prism.

### Mass photometry

Mass photometry was performed on a Refyn Two MP as previously described (71). Pol complexes were diluted from stocks to 50 nM in 50 mM Tris-HCl pH 7.5, 1 mM DTT, 5% glycerol (vol/vol) and 250 mM NaCl were incubated in the indicated amounts of H_2_O_2_ for 5 minutes at 30°C. Proteins were diluted to 5 nM in the 9 µL of the same buffer and deposited on the coverslip, and movies were acquired for 1 min. MW histograms were generated with the Refyn DiscoverMP software.

### Serial spot dilution assays

Yeast spot assays were performed with yeast harvested in exponential phase. Cells were serially diluted to 10_4_, 10_3_, 10_2_, and 10 cells/µL with sterile water; spots were 5µL.

### Western Blot

Cells lysis was performed according to ref (72). 2.5 ODs of yeast were pelleted 5 min at 1000xg and resuspended in 100 µL of water. 100 µL of 0.2M NaOH was added for 5 min at RT and spun for 30 sec at 5000xg. The pellet was resuspended in 60 µl of 2X sample prep buffer (250 mM Tris pH 6.8, 10% glycerol, 1% SDS, 10% β-mercaptoethanol, 0.005% bromophenol blue), boiled 3 min, and centrifuged 1 min at 21,000xg, collecting supernatant. 6 µl per lane was loaded on a 7.5% acrylamide gel for SDS-PAGE and transferred to nitrocellulose for western blotting. Membrane was blocked in 1% BSA in TBS (10 mM Tris pH 8.0, 150 mM NaCl). Antibody incubations and washes were in TBST (10 mM Tris pH 8.0, 150 mM NaCl, 0.1% Tween-20) with 1% BSA. Primary antibodies: rabbit anti-RNR1 (Agrisera #AS21 4608) at 1:3000, or mouse anti-α-tubulin (ABM # G094) at 1:8000 for a loading control. Secondary antibodies: DyLight goat anti-mouse 680 nm, or goat anti-rabbit 680 nm, at 1:15,000. Blots were detected on an Odyssey CLx infrared scanner.

## DISCUSSION

In this report, we stumbled on the fact that HU can arrest cells independent of the S-phase checkpoint and/or oxidative stress signaling via Mrc1. In trying to find the basis for this, we found that HU generates excessive nuclear ROS that, in addition to its known relationship in generating mutagenic oxidative DNA lesions like 8-oxo-guanine (73), is likely responsible for inhibiting the replicative polymerases required for S-phase progression, particularly at the high concentrations of HU (∼50-200 mM) used to arrest yeast cells: the direct relationship between HU and H_2_O_2_ implies a level of H_2_O_2_ that is not only well above the ∼0.1mM H_2_O_2_ that elicits the cellular oxidative stress response (62, 74) but also the ∼10 mM H_2_O_2_ that we observed to fully inhibit replication *in vitro*. In this study, we could only observe that Pol complexes containing Fe-S clusters (Pols α, δ, and ε) were inhibited by H_2_O_2_, while Klenow Pol was not. Because the active sites of eukaryotic and prokaryotic DNA polymerases are structurally and functionally homologous (81), we attribute the stark contrast observed between the H_2_O_2_ sensitivity with the existence of oxidation-sensitive Fe-S clusters in the eukaryotic Pol complexes that do not exist in prokaryotic Pols. Notably, the fact that both DNA binding and polymerase activity of Klenow Pol were unfazed by profuse H_2_O_2_ also rules out the possibility that oxidation of either the DNA substrate or the dNTPs in the reaction was responsible for inhibition of either DNA polymerase and/or substrate binding activities of Pol α, δ, or ε.

It is well established that HU inhibits RNR, however, given the fact that dNTP pools are not appreciably reduced in yeast (37), and that HU can inhibit archaea with another class of RNR that do not require a tyrosyl radical (36), we consider it unlikely that RNR inhibition is the primary mechanism of cell cycle arrest by HU, particularly at the levels of HU used in eukaryotes. Importantly, the only studies to our knowledge that have reported RNR inhibition to be the sole mechanism of HU action are from bacteria (75–78), where Fe-S clusters are not coordinated in polymerases, and where significantly less HU is required. Keeping this in mind, only one study to our knowledge has claimed that HU is not a direct inhibitor of DNA synthesis *in vivo*, and this study was also performed in *E. coli* (79). Instead, at the high concentrations of HU routinely employed in eukaryotic cells, we predict that polymerases are likely one of a multitude of classes of Fe-S containing enzymes that are inhibited in the strong oxidative environment seen with HU incubation, that cellular pathways related to redox sensing are likely also activated, and that the molecular mechanisms of cytotoxicity by HU are likely manyfold. Still, we note that in theory, inhibition of any of the replicative polymerases would be enough to stall the cell cycle. The fact that perturbation of CysB in Pol δ and CysX in Pol ε inhibits these critical polymerases *in vivo* (57, 59, 60), and similarly that primase activity relies on Fe-S coordination (48, 51) indicates that oxidization of the integral Fe-S cluster in these complexes would be sufficient to stall cells. Critically, it was shown that mutation of the Cys residues that coordinate Fe-S in the CTD of human Pri2 (the regulatory subunit of primase) results in defective Pol α loading and initiation, with a temperature sensitive phenotype of G1/S arrest (80). Thus, oxidation-driven inhibition of just one of the three enzyme complexes we observed in this study is in fact sufficient to cause cell cycle arrest.

In an attempt to elucidate the mechanism of Pol inhibition by H_2_O_2_, we found that both CysA and CysB were dispensable for Pol α activity, indicating that a Fe-S and/or Zn^2+^ cofactor is likely not required for the polymerase activity of Pol1. A recent report describes a possible mechanism in human Pol α primase whereby primase may hand off to the polymerase subunit via a charge transfer between two Fe-S clusters, implying that the CTD of the Pol1 subunit would need to have a reduced Fe-S cluster in its CTD order to facilitate primase-to-polymerase handoff (54), however we do not see evidence for coordination of a charge transfer donor in Pol1 by either CysA or CysB. Recently, O’Donnell, Li, and colleagues published a structure of yeast Pol α comprising all 4 subunits, where a Fe-S cluster was not observed in the Pol1 CTD, corroborating our results (82). Because we were able to abrogate Pol α activity *in vivo* by disrupting the CysA or CysB domains in Pol1 CTD, but were able to strongly inhibit template/primer binding and elongation of Pol α with H_2_O_2_ *in vitro*, we predict that most of the DNA binding affinity of Pol its DNA elongation mode might come from the primase regulatory subunit, Pri2, previously observed to possess Fe-S clusters in both yeast and humans (48–53) and to remain bound to the template during the DNA polymerization step (82). Still, given the conservation of the CysA and CysB motifs in Pol1, and the fact that Fe-S was previously observed to associate with the Pol1 CTD (55), further study is warranted to understand the functional role, if any, of these motifs in Pol α.

How exactly does the 4Fe-FS cluster modulate polymerase activity? Based on our studies and work from other labs (36, 55, 58) it appears that these clusters may play mostly a structural role in helping maintain polymerase complex assembly and/or function. The Fe-S cluster seemingly plays a different role in each polymerase: The dissociation of Pol α subunits results in reduced DNA binding affinity, possibly by the loss of Pri2 binding as discussed above, while in Pol δ, the loss of the small subunits (Pol31 and Pol32) would eliminate its affinity for PCNA sliding clamp, which is required for activity (55, 83). In this work, Pol ε dissociated in the presence of H_2_O_2_ but was more stable than Pol α or δ. It remains unclear whether Pol ε retains an Fe-S cluster in its CTD (46), however a separate Fe-S cluster target of H_2_O_2_ seems more likely: mutagenesis of the CysX motif of the Pol2 NTD, which has been observed in structures to coordinate a Fe-S cluster, resulted in loss of DNA binding and activity as consequence of disrupting the P-domain, part of the catalytic core shown to be critical for processivity (84, 56, 57). If Pol ε indeed only retains an Fe-S cluster in the Pol2 NTD, it would appear this location is much less critical for Pol complex stability than Pri2 for Pol α or the CTD of Pol3 for Pol δ. Thus, all three polymerases are vulnerable to oxidation but by separable mechanisms. It is possible that coordination of Fe-S clusters by polymerases evolved as a “kill switch” to prevent DNA from being replicated in a potentially damaging environment by triggering rapid complex disassembly and/or unfolding under oxidizing conditions. Interestingly, it is even possible that these cofactors are communicating their redox status through DNA via charge transfer signaling (85). It will take further structural and genetics studies to fully understand their role(s) in replication. Regardless of the precise mechanism of inhibition by HU, the abundant ROS that it generates in cells should be recognized as a major confounding variable to present and past studies in eukaryotes utilizing HU-induced cell cycle arrest. Here we demonstrate that RNR-deg is a superior alternative to HU that is significantly more specific against RNR and allows cells to recover from arrest faster than in HU, all in the absence of toxic ROS generation. We hope that RNR-deg can be a useful tool to investigate the S-phase checkpoint, genome maintenance, DNA damage repair, and countless other processes in the cell without the confounding presence of HU-induced oxidative stress.

## ACKNOWLEDGEMENTS

We thank the Colorado State University (CSU) Core Facility (RRID:SCR_022000) for training and assistance with flow cytometry, cell sorting, and single cell analysis. We thank Kyleigh Castillo (NSF REU) for assistance with the plate assay. We thank the Markus and DeLuca labs at CSU for use of their Mass Photometer. We thank the O’Donnell lab at Rockefeller University for generously sharing protein expression strains and plasmids. Funding was provided by NIGMS (R00GM126143 and R35GM147105 to G.D.S.)

