## Supplemental Information for "Revised Mechanism of Hydroxyurea Induced Cell Cycle Arrest and an Improved Alternative"

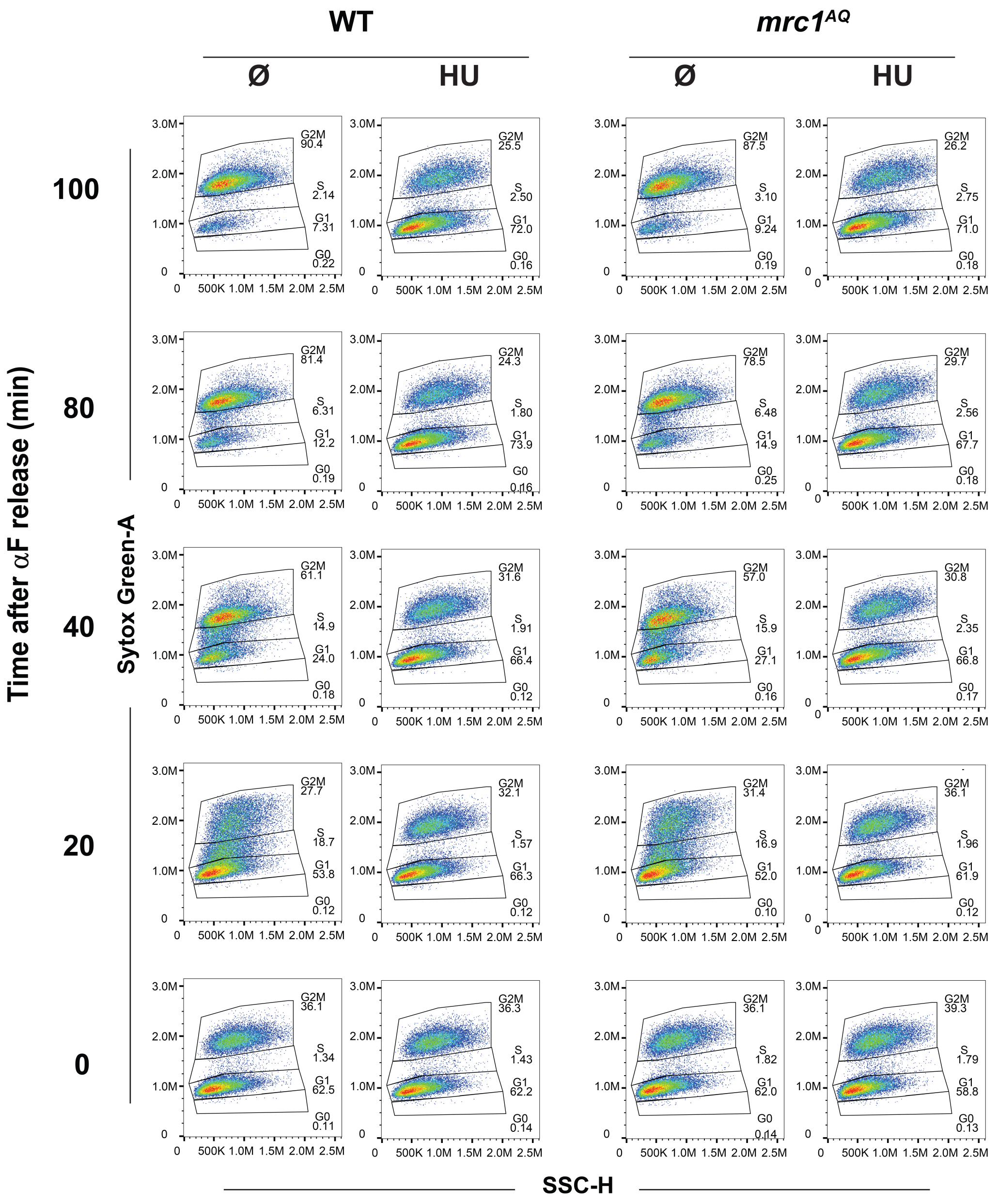


**Figure S1. Scatter plots from all experiments in Fig 1b.** Sytox Green vs. side scatter height plots for every panel in Fig 1b., including representative gating criteria used for study.

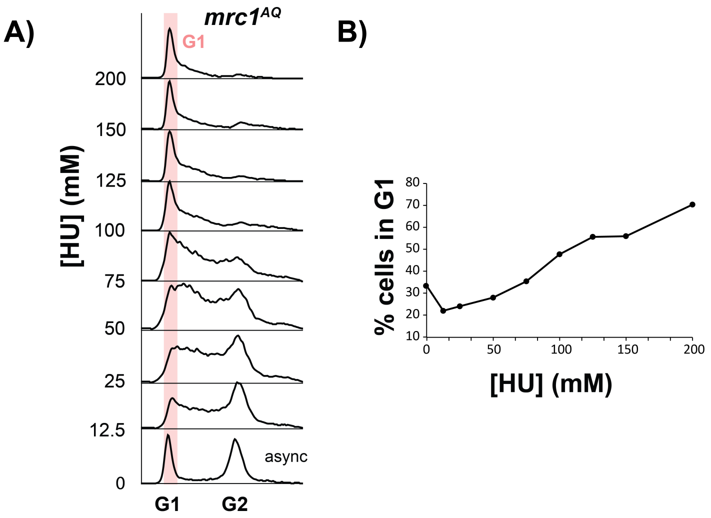


**Figure S2. HU arrests replication checkpoint deficient cells in G1/S at high concentrations. (A)** Cytometry profiles of *mrc1*$^{AQ}$ are shown at the indicated HU concentration, which was added to asynchronous cells growing in YPD for 4 hours prior to flow cytometry acquisition. Red bar indicates G1. **(B)** Plot of percentage cells in G1 from the same experiment.

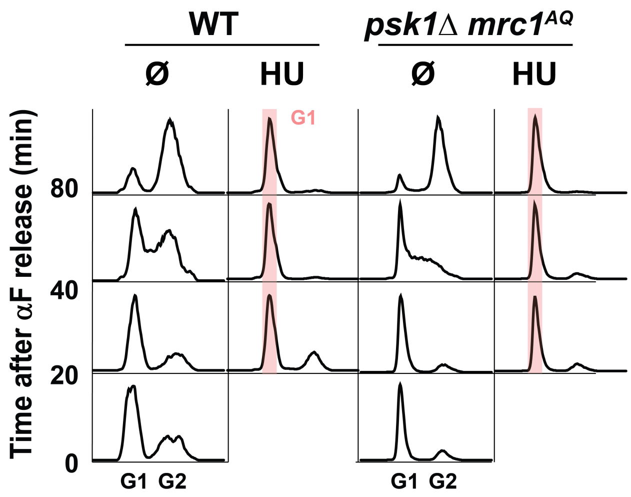


**Figure S3. HU arrests cells in G1/S in the absence of Psk1/Mrc1 and Mec1/Mrc1 signaling.** WT or *psk1Δ mrc1*$^{AQ}$ cells were synced in G1 with αF and released into YPD in the absence or presence of 0.2 M HU, and cytometry profiles were acquired at the indicated times. Red bar indicates G1/S stalling.

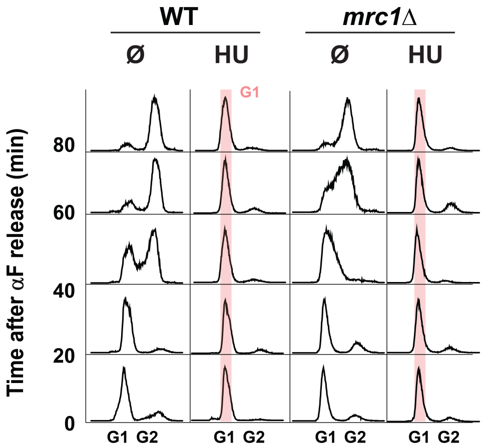


**Figure S4. HU arrests cells in G1/S in the absence of Mrc1.** WT or *mrc1Δ* cells were syncrhonized in G1 by αF arrest and released into YPD in the absence or presence of 200 mM HU, and cytometry profiles were acquired at the indicated times. Red bar indicates G1/S stalling.

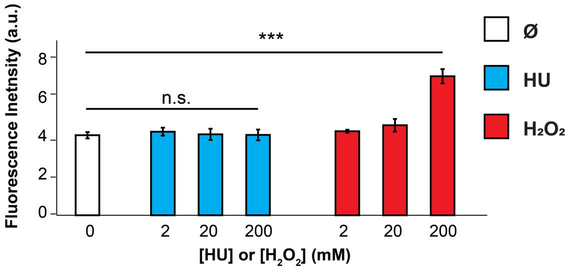


**Figure S5. HU does not oxidize or activate the ROS probe DHE.** *In vitro* reactions containing plasmid DNA, DHE, and the indicated amount of HU or $H_2O_2$ were carried out in a 384-well plate reader. Red-shifted fluorescence (610 nm Em) is shown; error bars represent STD of experiments repeated in triplicate. Single-factor ANOVA was used to test the difference between the control group and all HU concentrations (n.s.; p=0.63) or the control group and all $H_2O_2$ concentrations ($p=1.9x10^{-6}$).

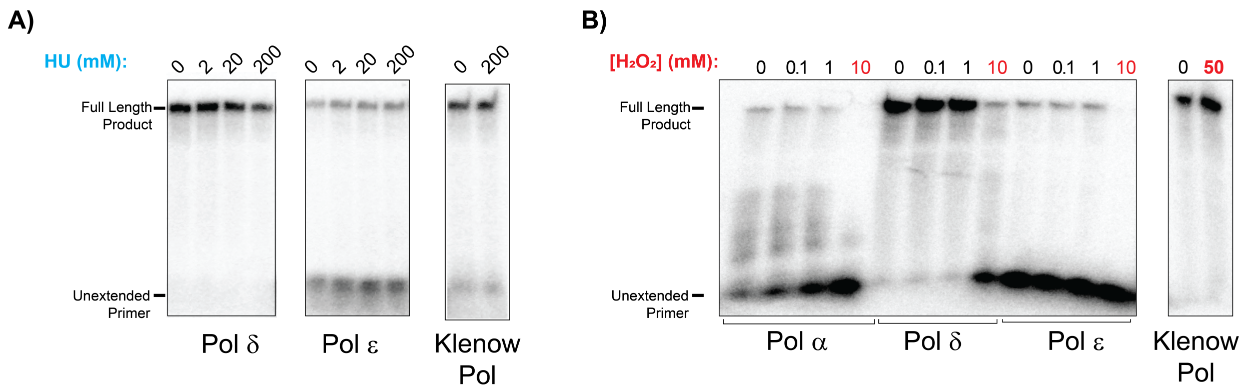


**Figure S6. Strand extension gels.** Denaturing PAGE gels show strand extension activity of indicated proteins in the presence of the indicated amounts of **(A)** HU or **(B)** $H_2O_2$ (see assay schematic in Fig 3a). The gels in (B) are representative of one of three separate experiments used to calculate replication inhibition bar graphs shown in Fig 3a.

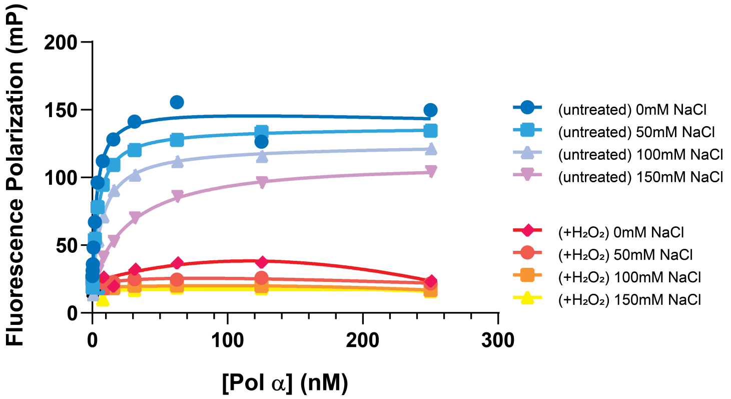


**Figure S7. Salt screening of Pol α binding to DNA template primer.** NaCl was titrated into Pol a:template/primer binding reactions at the indicated concentrations, in the absence or presence of 50 mM $H_2O_2$. Binding was determined by reading fluorescence polarization of FITC in a 384-well plate at the indicated Pol α concentrations.

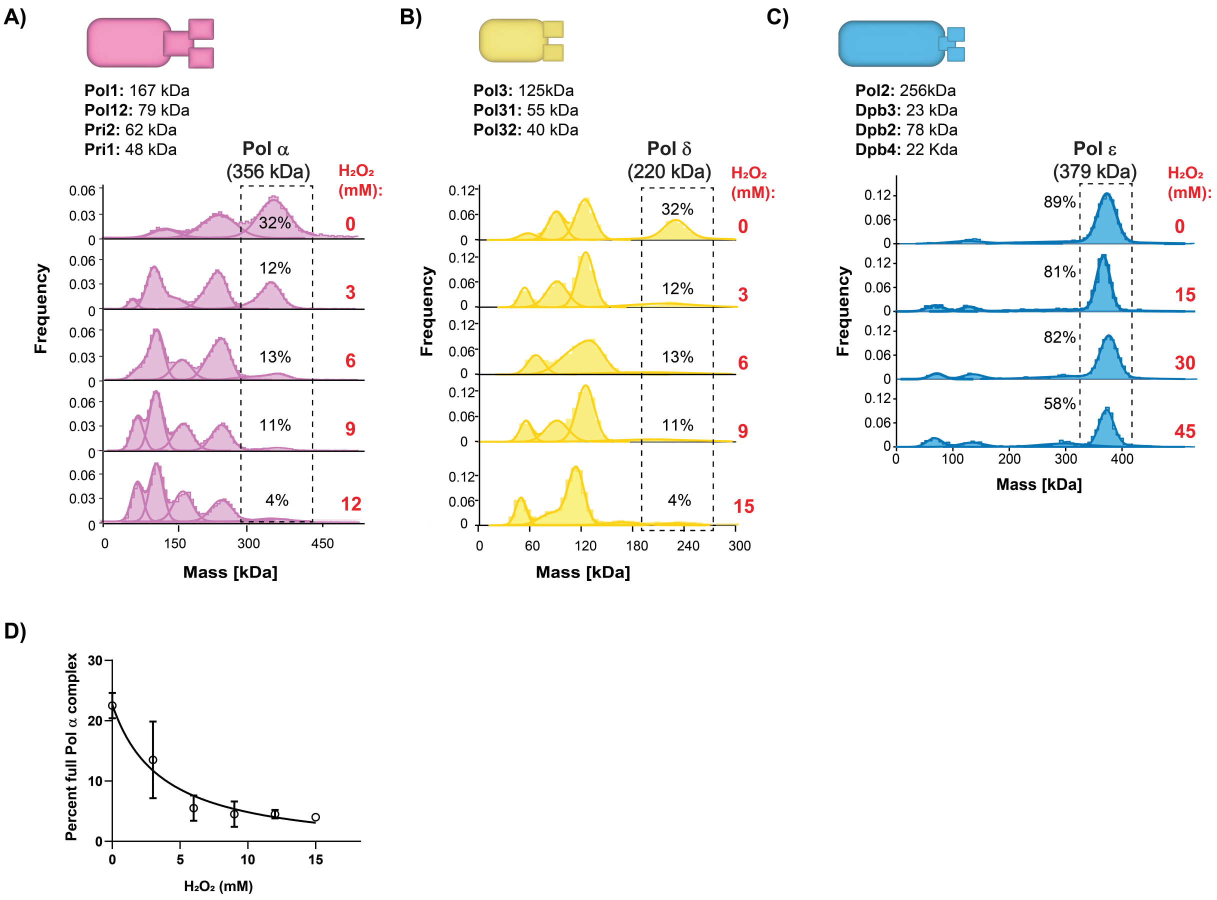


**Figure S8. Supplemental mass photometry data for Pols α, δ, and ε.** Mass photometry histograms for **(A)** Pol δ, **(B)**, Pol δ and **(C)** Pol ε are shown at the indicated $H_2O_2$ concentrations in red. A cartoon of each Pol is shown, and the MW of each subunit is listed. Full complex MW is denoted by dotted rectangle, and the relative percentage of that full complex is shown on each panel. **(D)** Titration curve of percent full Pol α complex vs [$H_2O_2$] is shown. Error bars represent STD from two separate mass photometry experiments.

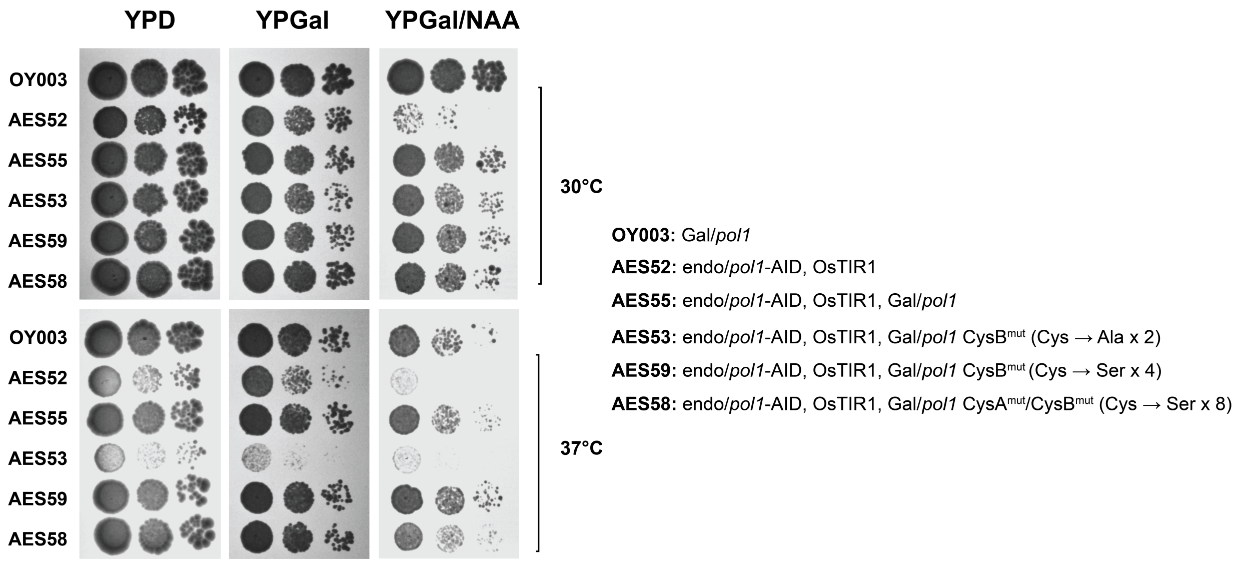


**Figure S9. CysA and CysB, the metal binding sites in the CTD of Pol1, are dispensable for Pol α activity *in vivo*.** Yeast spot survival assays are shown for the indicated strains, in the indicated conditions, at the indicated temperature. Spots contained 5000, 500, or 50 cells. "endo/*pol1*-AID" encodes an AID degron tag on the CTD of endogenous Pol1. "Gal/*pol1*" denotes an overexpressed copy of *pol1* at a different locus. Both OsTIR1 and, if present, Gal/*pol1*, are expressed in the presence of Gal. OsTIR1 requires NAA to degrade AID-tagged proteins. The positions of the CysA/B mutations are described in Supplementary Methods.

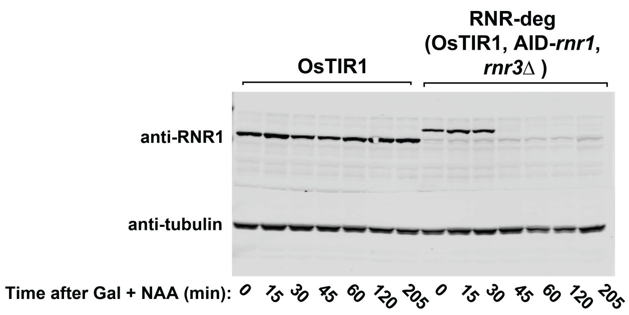


**Figure S10. Western blot time course of RNR-deg.** Cells from the indicated strains were treated with Gal+NAA, lysed at the indicated times, and separated by SDS-PAGE. The membrane was probed with a primary rabbit anti-Rnr1 followed by a secondary goat anti-rabbit 680 nm antibody. Rnr1-AID runs slower than Rnr1 due to its degron tag. Tubulin is tracked as a loading control (primary: mouse anti-a-tubulin; secondary: goat anti-mouse 680 nm). The gel was imaged with an infrared scanner.

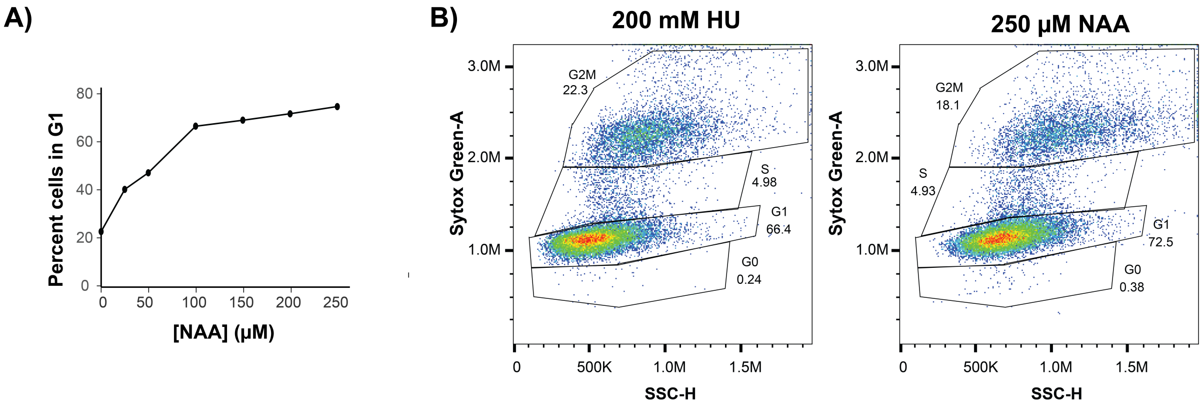


**Figure S11. Further RNR-deg characterization. (A)** Extent of cycle arrest in the RNR-deg strain is tunable by NAA concentration. Percentage of cells stalled in G1, quantified in cytometry data using identical gates, is plotted against [NAA] **(B)** Characterization of cell shape in RNR-deg vs. HU treatment. Sytox Green vs Side scatter height plots are shown after treatment of asynchronous RNR-deg cells (OsTIR1, AID-*rnr1*, *rnr3Δ*) with 200 mM HU or 250 μM NAA for 4 hours.

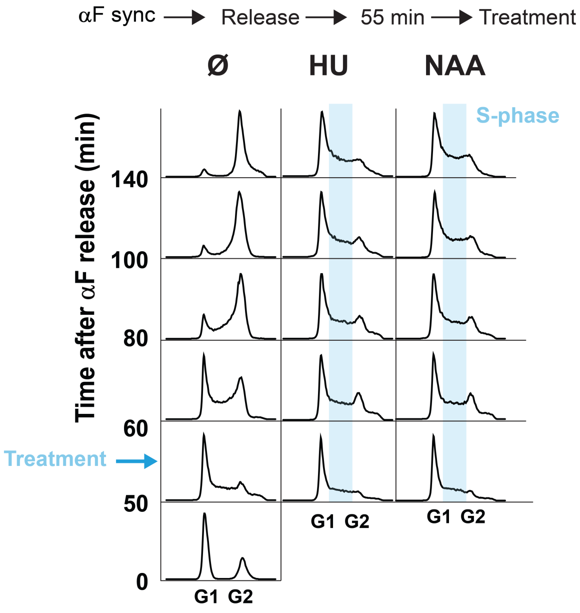


**Figure S12. Robust S-phase arrest with RNR-deg.** RNR-deg cells (OsTIR1, AID-*rnr1*, *rnr3Δ*) were synchronized in G1 with αF and released into YPG/Gal for 55 minutes, followed by treatment with either HU or NAA (see schematic). Flow cytometry profiles were acquired at the indicated times. The blue bar indicates S-phase.

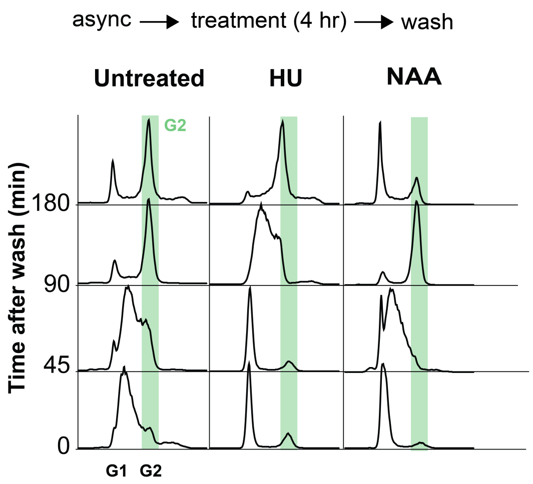


**Figure S13. RNR-deg treatment reverses faster than HU treatment after washout.** Asynchronous RNR-deg cells were treated with $\alpha$F, released, released into media containing the indicated treatment for 4 hours, and washed extensively by pelleting (1000xg) then resuspending cells in water two times, followed by a final pelleting/resuspension in YPG/Gal. Cytometry profiles were acquired at the indicated times after washout. Green bar highlights cells in G2. The cell cycle speed of RNR-deg + NAA cells after wash is faster than post-wash HU-treated cells, comparable to the speed of untreated cells.

### SI MATERIALS AND METHODS

**RNR-deg construction.** Gal controlled OsTIR1 was integrated into the *URA3* gene of OY01 (*ade2-1 ura3-1 his3-11,15 trp1-1 leu2-3,112 can1-100 bar1Δ MATa pep4::KANMX6)* using the GAL-OsTIR1(WT) plasmid from Addgene (cat # 140655) (86) to create AES25. A miniAID1 tag was added to the N-terminal of endogenous *RNR1* using CRISPR/Cas9, following previously detailed protocols (87). Q5 mutagenesis (NEB) was used to modify Addgene gRNA plasmid #64330 (88) to target RNR1 (target: ATGTACGTTTATAAAAGAGA). *RNR3* was replaced by a HygR cassette. A gene block donor (Twist Biosciences) of AID-*rnr1* was amplified by PCR. 3 µg of this donor DNA, along with 0.7 µg each of cas9 (#64329, nourseothricin selection) and *RNR1* gRNA plasmids (hygromycin selection) was transformed into AES25 with selection on 100 µg/ml NTC and 700 µg/ml HygB. The resulting strain was AES30 (OsTIR1, AID-*rnr1*). Finally, RNR3 was knocked out of AES30 and replaced with a hygromycin resistance cassette. The strains had previously been tested to ensure that they had lost the transiently transformed gRNA plasmid which carried hygromycin resistance: a donor DNA was created that amplified the hygromycin resistance cassette from Addgene plasmid #64330 and added 69 bp of homology each to *RNR3* 5' and 3' UTRs. 6 µg of donor DNA was transformed, with selection on 700 µg/ml hygromycin B. Colonies were screen by PCR for successfully integrated HygR cassette. The resulting strain was AES38 (RNR-deg); (OsTIR1, AID-*rnr1, rnr3Δ*).

***mrc1Δ*, *mrc1$^{AQ}$*, *mrc1$^{AQ}$ psk1Δ*.** *mrc1* knockout was performed by replacement with a *URA3* (AT07) or a *TRP1* (AT03). *mrc1$^{AQ}$* (AT04) was created by integrating *mrc1$^{AQ}$* donor into the *mrc1Δ* locus of AT03 and counterselecting with **5-fluoroanthranilic acid (5-FAA)**. A *psk1* knockout was performed by replacing *psk1* with the **nourseothricin N-acetyltransferase (NAT)** expression cassette from Addgene plasmid #64329 (83) to create *mrc1$^{AQ}$ psk1Δ* (AES44).

**Pol1-AID:** An AID tag was added to the C-terminus of endogenous Pol1 in AES25 (OsTIR1) or OY003, a strain containing an integrated copy of *GAL-pol1*-3xFlag as previously described (65) by targeting the 3'UTR of the *POL1* gene with CRISPR/cas9. The target in both strains was CTAATGTACAGACATCGGA, which is not present in the 3'UTR of the pol1-3xFlag in OY003. This resulted in AES52 (*pol1*-AID, OsTIR1). OsTIR1 was moved into pRS405/GAL (67) and integrated into the LEU2 marker of the (*pol1*-AID, *GAL-pol1*-3xFlag) strain to make AES55 (Pol1-AID, OsTIR1, GAL-Pol1).

**Pol1 CysA and CysB mutants:** The potential Fe-S motifs of Pol1 (55) were mutated in the Gal1 controlled, exogenous copy of *pol1*-3xFlag in AES55, the *pol1*-AID,OsTIR1,Gal/Pol α strain described in the previous step. Three versions were made mutating the possible Fe-S motifs:

CysB (AES53) is C1353A,C1372A (two critical Cys from CysB)

CysAB (AES59) is C1287S,C1290S,C1314S,C1317S (CysA) and C1348S,C1353S,C1367S,C1372S (CysB)

CysBx4 (AES58) is C1348S,C1353S,C1367S,C1372S (CsyB)

These mutations were introduced using CRISPR/Cas9, by targeting a gRNA to the flag tag of *pol1*-3xFlag (target: GGATGACGATGACAAGTGAG). The donors for homologous recombination had extensive homology to the 3'UTR specific to *pol1*-3xFlag to avoid targeting the endogenous copy of Pol1. The donors were silently mutated to introduce a PmeI restriction site to facilitate screening of modified clones.

**DNA strand extension template construction.** The primed substrate used to test the polymerase activity of Pol α, δ, and ε required assembly of the leading strand was described previously (21). To create the 243 nt leading strand, Near_143 and Far_100 were ligated together by first mixing with Near_Far_bridge in a 1:1:3 molar ratio in the presence of 50 mM NaCl and 5 mM trisodium citrate pH 7.0, followed by annealing by heating to 94°C for 5 min and allowing to cool slowly to room temperature.

Following this, 4,000 Units of T4 Ligase and 1x T4 ligase buffer (New England Biolabs) and an additional 1 mM ATP were added, and the reaction was incubated for 16 h at 15°C. The ligated product was purified on an 8% denaturing PAGE gel using SYBR Safe stain (Invitrogen) to visualize and excise the single-stranded product. The DNA was recovered by crushing the gel and soaking in buffer TE pH 8.0 for 16 h at room temperature, followed by spinning at 15,000 rpm for 5 min, and aspirating the supernatant. The resultant 243 nt leading strand oligo was annealed to 85DS_primer, was 5'-end labeled with $\gamma$-$^{32}$P-ATP (Revvity) by T4 PNK (New England Biolabs) according to manufacturer instructions and purified on an S-200 HR microspin column (GE Healthcare). The radiolabeled primer was annealed to the fork at a 1:1 ratio.

**SI Table 1: Yeast strains used in this study. All strains are derived from W303 (*S.Cerevisiae*).**

| Strain Name | Genotype |
|---|---|
| OY01 | *ade2-1 ura3-1 his3-11,15 trp1-1 leu2-3,112 can1-100 bar1Δ MATa pep4::KANMX6* (from ref 65) |
| AES25 | *ade2-1 ura3-1::GAL-OsTIR1 his3-11,15 trp1-1 leu2-3,112 can1-100 bar1Δ MATa pep4::KANMX6* |
| AES30 | *ade2-1 ura3-1:: GAL-OsTIR1 his3-11,15 trp1-1 leu2-3,112 can1-100 bar1Δ MATa pep4::KANMX6 AID-rnr1* |
| AES37 | *ade2-1 ura3-1:: GAL-OsTIR1 his3-11,15 trp1-1 leu2-3,112 can1-100 bar1Δ MATa pep4::KANMX6 AID-rnr2* |
| AES38 | *ade2-1 ura3-1:: GAL-OsTIR1 his3-11,15 trp1-1 leu2-3,112 can1-100 bar1Δ MATa pep4::KANMX6 AID-rnr1 rnr3-Δ::HygR* |
| AT07 | *ade2-1 ura3-1 his3-11,15 trp1-1 leu2-3,112 can1-100 bar1Δ MATa pep4::KANMX6 mrc1-Δ::URA3* |
| AT03 | *ade2-1 ura3-1 his3-11,15 trp1-1 leu2-3,112 can1-100 bar1Δ MATa pep4::KANMX6 mrc1-Δ::TRP1* |
| AT04 | *ade2-1 ura3-1 his3-11,15 trp1-1 leu2-3,112 can1-100 bar1Δ MATa pep4::KANMX6 mrc1-Δ::mrc1$^{AQ}$* |
| AT02 | *ade2-1::GAL-mrc1$^{AQ}$-3xFlag ura3-1 his3-11,15::GAL-tof1,15 trp1-1 leu2-3::GAL-Csm3-6xHis,112 can1-100 bar1Δ MATa pep4::KANMX6* |
| AES44 | *ade2-1 ura3-1:: GAL-OsTIR1 his3-11,15 trp1-1 leu2-3,112 can1-100 bar1Δ MATa pep4::KANMX6 mrc1-Δ::mrc1$^{AQ}$ psk1-Δ::NAT* |
| OY003 | *ade2-1::GAL-pol1-3xFlag ura3-1 his3-11,15 trp1-1 leu2-3,112 can1-100 bar1Δ MATa pep4::KANMX6* (from ref 64) |
| AES52 | *ade2-1 ura3-1::OsTIR1 his3-11,15 trp1-1 leu2-3,112 can1-100 bar1Δ MATa pep4::KANMX6 pol1-AID* |
| AES55 | *ade2-1::GAL-pol1-3xFlag ura3-1 his3-11,15 trp1-1 leu2-3,112::OsTIR1 can1-100 bar1Δ MATa pep4::KANMX6 pol1-AID* |
| AES53 | *ade2-1::GAL-pol1(C1353A,C1372A)-3xFlag ura3-1 his3-11,15 trp1-1 leu2-3,112::OsTIR1 can1-100 bar1Δ MATa pep4::KANMX6 pol1-AID* |
| AES59 | *ade2-1::GAL-pol1(C1287S,C1290S,C1314S,C1317S, C1348S,C1353S,C1367S,C1372S)-3xFlag ura3-1 his3-11,15,15 trp1-1 leu2-3,112::OsTIR1 can1-100 bar1Δ MATa pep4::KANMX6 pol1-AID* |
| AES58 | *ade2-1::GAL-pol1(C1348S,C1353S,C1367S,C1372S)-3xFlag ura3-1 his3-11,15 trp1-1 leu2-3,112::OsTIR1 can1-100 bar1Δ MATa pep4::KANMX6 pol1-AID* |

**SI Table 2. DNA oligonucleotides used in this study.** Sequences up to 200 nt were synthesized by IDT, and gene blocks were synthesized by Twist Biosciences.

| Name | Sequence |
|---|---|
| AID-RNR1 donor | CTCTACCATAATTGAAGCATATCTCATCCTTTTCATCCTTTTCAACGCAAGAGAGACACCAACGAACAACACTTTATTTGTTGATATATTAACATCAAAAAATGTCTAAAGAAAAATCTGCTTGTCCAAAAGATCCTGCTAAACCACCTGCTAAAGCTCAAGTTGTTGGTTGGCCACCTGTTAGATCTTATAGAAAAAATGTTATGGTTTCTTGTCAAAAATCTTCTGGTGGTCCTGAAGCTGCTGCTTTTGTTAAAGTTTCTATGGATGGTGCTCCATATTTGAGAAAAATTGATTTGAGAATGTATAAAGCTTCTATGTATGTCTACAAGCGTGACGGTCGTAAAGAACCTGTCCAATTCGATAAGATTACCGCTCGTATATCACGCTTATGCTATGGTTTAGATCCAAAACATATCGACGCCGTTAAGGTCACCCAACGTATCATTTCTGGTGTCTAT |
| RNR3_HygR_fwd | TCATATCCAAGTTGAAATAAATATGACAAGCAAGAATAGCAGCAGCAATAAATCAAATACTCCCACACAGACATGGAGGCCCAGAATACC |
| RNR3_HygR_rev | TAATACATACTAACGAAAAGAAACCGCTCCAAGTTAGATAAGGAAAGGGAAAAATGCCACCAGAAAGAATTATTCCTTTGCCCTCGGACG |
| PSK1_NAT_fwd | CTATTTTTGTTTTGTTTTGTTTTGTTTTGTTCTCTCCCTAATTTGCATATAGGTAAACATCAAAGAAGTGACATGGAGGCCCAGAATACC |
| PSK1_NAT_rev | CATTGTTTCTACTGGATATTTTTTTGCATTTTCTTTTCCAATTACTAATGTCCTAGAATGATCATTAATCCTTTAGGGGCAGGGCATGC |
| Pol1-AID_donor | GCTTTAACAGAACAAAACAGAGAACTAATGGAAACCGGCCGGAGCGTTGTTCAAAAATATTTGAACGATTGTGGACGTCGCTACGTTGATATGACTAGCATATTTGATTTCATGCTAAATGGTGGTTCTAAAGAAAAATCTGCTTGTCCAAAAGATCCTGCTAAACCACCTGCTAAAGCTCAAGTTGTTGGTTGGCCACCTGTTAGATCTTATAGAAAAAATGTTATGGTTTCTTGTCAAAAATCTTCTGGTGGTCCTGAAGCTGCTGCTTTTGTTAAAGTTTCTATGGATGGTGCTCCATATTTGAGAAAAATTGATTTGAGAATGTATAAATAGCCTTATGTACTGACATCGCAACGAGCGCTAGGAAGCTTTTATTATTGAAGGTTTTCAGCGATATTCTGAGAATGTTTTTTCGGCTCTGGTGATCAGTTCTCACATGCATCATCCTTTAATAAACCCATGCATTAAGATTCGCC |
| CysB_donor | GGATACTGTAACATTAGAATTAAGTTGCCCATCATGCGATAAAAGGTTTCCATTTGGTGGTATTGTATCTTCAAATTACTATCGCGTGTCATATAATGGTTTACAGTGCAAGCATTGTGAGCAACTTTTTACTCCTCTTCAATTAACTAGCCAAATAGAGCATTCTATAAGGGCACACATTTCCTTATATTACGCAGGGTGGTTACAGTGTGATGACAGCACAGCTGGTATAGTTACAAGACAAGTCTCCGTTTTTGGTAAGCGTTGTTTAAACGACGGCGCTACGGGTGTCATGAGATACAAATACAGTGACAAGCAATTGTACAATCAACTTTTGTATTTCGATTCTTTGTTCGATTGTGAGAAGAACAAAAAGCAAGAATTGAAGCCAATATATCTACCCGATGATCTCGACTACCCCAAGGAACAGCTGACAGAATCATCTATTAAGGCTTTAACAGAACAAAACAGAGAACTAATGGAAACCGGCCGGAGCGTTGTTCAAAAATATTTGAACGATTGTGGACGTCGCTACGTTGATATGACTAGCATATTTGATTTCATGCTAAACGATTATAAGGATCATGACGGTGATTATAAAGATCATGACATCGACTACAAGGATGACGATGACAAGTGAGCGGCCGCCACCGCGGTGGAGCTCCAGCTTTTGTTCCCTTTAGTGAGGGTTAATTGCGCGCTTGGCGTAATCATGGTCATAGCTGTTTCCTGTGTGAAATTGTTATCCGCTCACAATTCCACACAACATACGAGCCGGAAGCATAAAGTGTAAAGCCTGGGGTGCCTAATGAGTGAGCTAACTCACATTAATTGCGTTGCGCTCACTGCCCGCTTTCCAGTCGGGAAACCTGTCGTGCCAGCT |
| CysAB_donor | GGATACTGTAACATTAGAATTAAGTTGCCCATCATGCGATAAAAGGTTTCCATTTGGTGGTATTGTATCTTCAAATTACTATCGCGTGTCATATAATGGTTTACAGTGCAAGCATTGTGAGCAACTTTTTACTCCTCTTCAATTAACTAGCCAAATAGAGCATTCTATAAGGGCACACATTTCCTTATATTACGCA |

| | |
|---|---|
| | GGGTGGTTACAGTCTGATGACAGCACATCTGGTATAGTTACAAGACAA GTCTCCGTTTTTGGTAAGCGTAGTTTAAACGACGGCTCTACGGGTGTC ATGAGATACAAATACAGTGACAAGCAATTGTACAATCAACTTTTGTATT TCGATTCTTTGTTCGATTGTGAGAAGAACAAAAAGCAAGAATTGAAGCC AATATATCTACCCGATGATCTCGACTACCCCAAGGAACAGCTGACAGA ATCATCTATTAAGGCTTTAACAGAACAAAACAGAGAACTAATGGAAACC GGCCGGAGCGTTGTTCAAAAATATTTGAACGATTGTGGACGTCGCTAC GTTGATATGACTAGCATATTTGATTTCATGCTAAACGATTATAAGGATC ATGACGGTGATTATAAAGATCATGACATCGACTACAAAGACGATGACG ATAAGTAAGCCGCCGCCACCGCGGTGGAGCTCCAGCTTTTGTTCCCTT TAGTGAGGGTTAATTGCGCGCTTGGCGTAATCATGGTCATAGCTGTTT CCTGTGTGAAATTGTTATCCGCTCACAATTCCACACAACATACGAGCC GGAAGCATAAAGTGTAAAGCCTGGGGTGCCTAATGAGTGAGCTAACTC ACATTAATTGCGTTGCGCTCACTGCCCGCTTTCCAGTCGGGAAACCTG TCGTGCCAGCT |
| CysBx4_donor | GTGGTGCGTTTGAGTGAAGCGCTTGGTTTAGATAGTAAAAAGTATTTTA GAAGAGAGGGCGGTAATAACAATGGAGAAGATATTAATAATTTGCAAC CCTTAGAAACAACAATTACAGACGTTGAAAGGTTTAAGGATACTGTAAC ATTAGAATTAAGTTCCCCATCATCCGATAAAAGGTTTCCATTTGGTGGT ATTGTATCTTCAAATTACTATCGCGTGTCATATAATGGTTTACAGTCCAA GCATTCTGAGCAACTTTTTACTCCTCTTCAATTAACTAGCCAAATAGAG CATTCTATAAGGGCACACATTTCCTTATATTACGCAGGGTGGTTACAGT CTGATGACAGCACATCTGGTATAGTTACAAGACAAGTCTCCGTTTTTG GTAAGCGTAGTTTAAACGACGGCTCTACGGGTGTCATGAGATACAAAT ACAGTGACAAGCAATTGTACAATCAACTTTTGTATTTCGATTCTTTGTTC GATTGTGAGAAGAACAAAAAGCAAGAATTGAAGCCAATATATCTACCC GATGATCTCGACTACCCCAAGGAACAGCTGACAGAATCATCTATTAAG GCTTTAACAGAACAAAACAGAGAACTAATGGAAACCGGCCGGAGCGTT GTTCAAAAATATTTGAACGATTGTGGACGTCGCTACGTTGATATGACTA GCATATTTGATTTCATGCTAAACGATTATAAGGATCATGACGGTGATTA TAAAGATCATGACATCGACTACAAAGACGATGACGATAAGTAAGCCGC CGCCACCGCGGTGGAGCTCCAGCTTTTGTTCCCTTTAGTGAGGGTTAA TTGCGCGCTTGGCGTAATCATGGTCATAGCTGTTTCCTGTGTGAAATT GTTATCCGCTCACAATTCCACACAACATACGAGCCGGAAGCATAAAGT GTAAAGCCTGGGGTGCCTAATGAGTGAGCTAACTCACATTAATTGCGT TGCGCTCACTGCCCGCTTTCCAGTCGGGAAACCTGTCGTGCCAGCT |
| Near_143 | /5phos/ATATTTATAATTAATTAAATATAATTTTTTTTTTTTTTTTTTTTTTT TTTTTTTTTTTTTTTTTGAGGAAAGAATGTTGGTGAGGGTTGGGAAGTG GAAGGATGGGCTTTTTTTTTTTTTTTTTTTTTTTTTTTTTTTTTTT |
| Far_100 | AGAGAGTAGAGTGAGTTGTGGATGTGTAGAGTTGTTGTAGGAGAAGA GTTGTGAAGTGTGGAGTGAGAGAAGAGAAGAGAGAGTGATATATTAAT ATTAT |
| 85DS_primer | AGCCCATCCTTCCACTTCCCAACCCTCACC |
| template_55 | TCAAAGTATCCAAATGAACGAATATGACTTTAGGAGTAGAGCGCGTGA GGATACA |
| FITC_primer_dd | /56-FAM/TGTATCCTCACGGCGTCTA/3ddC/ |
